# Pancreatic injury induces β−cell regeneration in axolotl

**DOI:** 10.1101/2025.01.23.634564

**Authors:** Connor J. Powell, Hani D. Singer, Ashley R. Juarez, Ryan T. Kim, Duygu Payzin-Dogru, Aaron M. Savage, Noah J. Lopez, Steven J. Blair, Adnan Abouelela, Anita Dittrich, Stuart G. Akeson, Miten Jain, Jessica L. Whited

## Abstract

**Background:** Diabetes is a condition characterized by a loss of pancreatic β−cell function which results in the dysregulation of insulin homeostasis. Using a partial pancreatectomy model in axolotl, we aimed to observe the pancreatic response to injury.

**Results:** Here we show a comprehensive histological assessment of pancreatic islets in axolotl. Following pancreatic injury, no apparent blastemal structure was observed. We found a significant, organ-wide increase in cellular proliferation post-resection in the pancreas compared to sham-operated controls. This proliferative response was most robust at the site of injury. We found that β−cells actively contributed to the increased rates of proliferation upon injury. β−cell proliferation manifested in increased β−cell mass in injured tissue at two weeks post injury. At four weeks post injury, we found organ-wide proliferation to be extinguished while proliferation at the injury site persisted, corresponding to pancreatic tissue recovery. Similarly, total β−cell mass was comparable to sham after four weeks.

**Conclusions:** Our findings suggest a non-blastema-mediated regeneration process takes place in the pancreas, by which pancreatic resection induces whole-organ β−cell proliferation without the formation of a blastemal structure. This process is analogous to other models of compensatory growth in axolotl, including liver regeneration.

## Background

Diabetes is a condition characterized by a loss of pancreatic β−cell function which results in the body’s inability to maintain insulin homeostasis. Treatment for diabetes can be, in most cases, lifelong, requiring administration of exogenous insulin or glycemic control agents, and glucose monitoring. Diabetes can cause complications such as heart failure, kidney disease, and tissue necrosis requiring amputation.^1,2,3^ Several treatment strategies, such as cadaveric β−cell transplantation, polymer encapsulated stem cell-derived β−cell transplantation, transplantation of chemically induced pluripotent stem-cell-derived islets, and personalized endoderm stem cell-derived islet tissue, aimed at functional restoration, have been implemented to varying degrees of success.^4,5,6,7^ These strategies, however, are invasive procedures that do not address the root cause of initial β−cell loss of function. A thorough understanding of genetic pathways that can promote pancreatic regeneration is important for developing therapeutic strategies for diabetes based on stimulating endogenous β-cell replacement.

Pancreatic regeneration has been reported in several animal models, such as mice and zebrafish, with strong conservation in injury responses.^8,9^ Pancreatic regeneration studies typically focus on the endocrine pancreas i.e. β−cells due to their relevance to diabetes. β−cells are the principal insulin producing cells in the body, and characterizing their development, function, and fate has been the topic of research for decades. Partial pancreatectomy studies in rats and mice have reported proliferation of existing islet and exocrine cells, demonstrating substantial regenerative ability in these organisms.^8^ Further studies in mice demonstrate β−cell regeneration to be a result of pre-existing β−cell differentiation in conjunction with transdifferentiation of other endocrine cell types.^4,10^ Despite strong characterization of the pancreas in mammals, endogenous β−cell regeneration has not been accomplished in human patients.

The axolotl (*Ambystoma mexicanum*) is a highly regenerative species of salamander that can provide insights into how molecular mechanisms of regeneration might be harnessed therapeutically in humans. While the axolotl has been shown to regenerate several tissue types such as limbs, lungs, and liver; few studies have investigated pancreatic regenerative ability in axolotls.^11,12,13^ Conservation of pancreatic regeneration in many other organisms leads us to reason the axolotl may display a similar proficiency for pancreatic response to injury.

Although the pancreas is well studied in humans and mice, only one study has reported on the pancreas of axolotls.^14^ This study, which established a diabetic axolotl model via chemical ablation of β−cells using streptozotocin, demonstrated a return to normal glycemic function after 84 days. These results indicated a restoration of β−cell function – possibly due to regeneration. However, streptozotocin was reported to cause side effects such as edema, lowered red blood cell count, and increased mortality.^14^ These findings introduced the axolotl as a promising animal model to study diabetes and illustrated the need to develop a more consistent injury model to show whether axolotls exhibit robust pancreatic regeneration. We sought to expand on the limited literature currently available in regard to axolotl pancreas regeneration by studying the regenerative response to partial pancreatectomy in axolotls.

Here, we characterize pancreatic tissue morphology and the pancreatic response to injury in axolotls. We established a novel pancreatic resection surgery model to investigate how animals respond to this injury over the course of 28 days. We identified key genes such as *Pdx1*, *Ins*, *Tgf-β1, Ctrb2*, *Marco*, *Kazald2*, and *Cirbp* to be differentially regulated in response to pancreatic injury.

## Results

### A novel pancreatic resection model in the axolotl

We established a pancreatic resection surgery model with a sham-operated counterpart in order to investigate the injury response after a loss of pancreatic tissue mass (Figure 1A-B). Pancreatic resection was performed by surgical incision at a site adjacent to the base of the duodenum, resulting in removal of approximately twenty percent of the length of the organ. Animals with larger portions of their pancreas resected generally did not survive. Injury response was investigated at 14 and 28 days post injury (dpi). Ninety percent of animals survived the surgery, and death was non-specific to the sham or resection group. Mortality could have resulted from blood loss, infection, or other unknown causes. By 14 dpi, the abdominal incision wound had partially healed, while the pancreas appeared to have fully healed. The abdominal incision had fully healed by 28 dpi. No visible increase in tissue mass or blastema-like structure were observed at the site of pancreatic resection. Pancreatic resection studies in mice suggest that regeneration in pancreatic tissue can materialize in several forms, leading us to hypothesize that non-blastemal regeneration mechanisms such as transdifferentiation, endocrine cell-specific proliferation, or compensatory growth are possible and could have been utilized by the axolotl in response to injury.^15^

**Figure 1.**
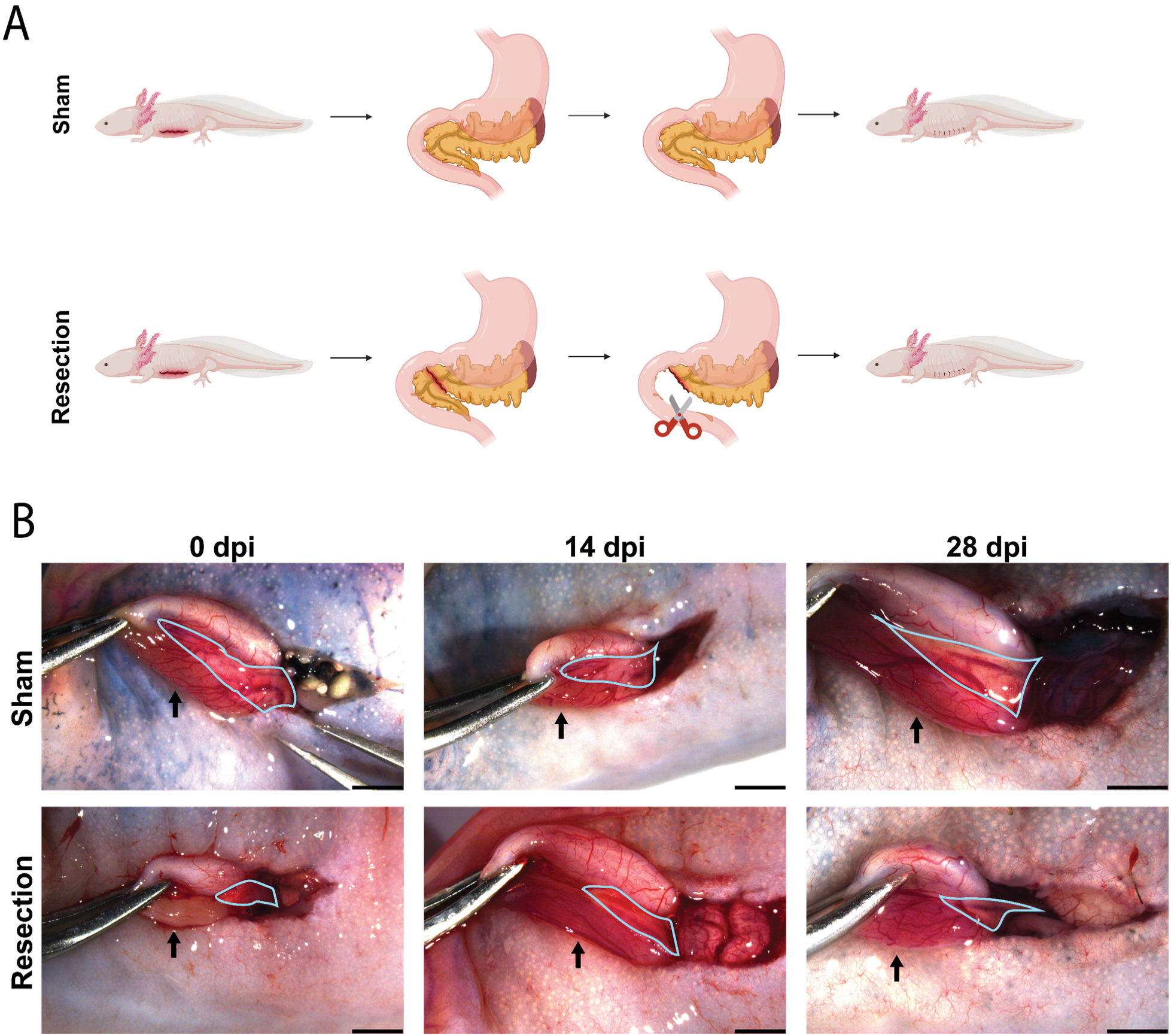
Pancreatic resection surgery model. **A.** Cartoon depicting sham and pancreatic resection surgery. **B.** Morphology of sham and pancreatic resection surgery at 0, 14 and 28 dpi. The pancreas is outlined in blue while black arrows point to the duodenum. 0 dpi image of the resection condition was taken immediately following 20% removal of pancreas by length. The majority of the pancreas extends into the abdominal cavity where it connects to the liver. In each image, the orientation from left to right follows a rostral to caudal direction. Black scale bars in the bottom right corner of each image are 2 mm.

### Pancreatic resection leads to significant increases in cellular proliferation

To assess cellular proliferation, we used 5-ethynyl-2’-deoxyuridine (EdU) staining on coronal sections of pancreas (whole organ) from the sham and resection animal groups. We used a single pulse of EdU, followed by a short chase period, to identify all cells which have recently gone through S-phase of the cell cycle. Organ-wide proliferation was analyzed and compared to proliferation within a standardized area local to the site of injury (Figure 2I). We found a significant increase in organ-wide cellular proliferation at 14 dpi in resected pancreas samples compared to the sham samples (Figure 2A−B). This proliferative response was significantly increased local to the site of injury in comparison to organ-wide proliferation (Figure 2C). Additionally, proliferation local to the site of injury was significantly increased in comparison to an analogous area in sham samples (Figure 2D, 2I). These results suggest that at 14 dpi, the axolotl pancreas undergoes both an organ-wide proliferative response to resection and a proliferative response local to the site of injury.

**Figure 2.**
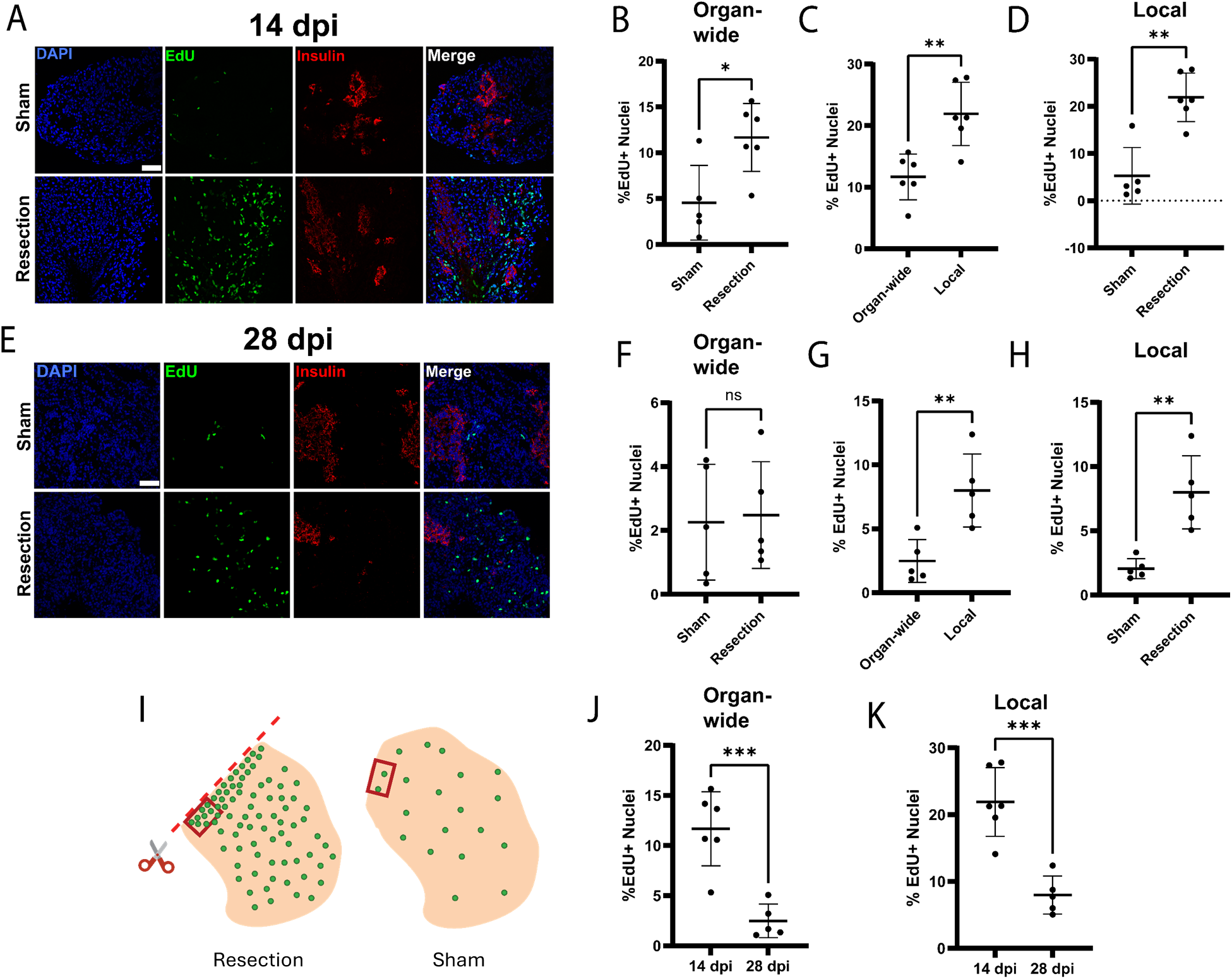
Pancreatic resection initially provokes substantial proliferative responses throughout the entire organ, but cell proliferation becomes restricted local to the site of injury later in the process. **A.** EdU and insulin (IHC) stain of pancreas samples at 14 dpi. **B.** Organ-wide proliferation is increased in resection samples at 14 dpi (p=0.0162) (sham n=5, resection n=6). **C.** Proliferation local to the site of injury is higher within resection samples at 14 dpi (p=0.0032). **D.** Proliferation local to the site of injury is increased at 14 dpi (p=0.0012). **E.** EdU and insulin stain of pancreas samples at 28 dpi (sham n=5, resection n=5). **F.** Organ-wide proliferation in resection samples is non-significant in comparison to sham at 28 dpi (p=0.8459). **G.** Proliferation local to the site of injury is higher within resection samples at 28 dpi (p=0.0085). **H.** Proliferation local to the site of injury is increased at 28 dpi (p=0.0079). **I.** Cartoon representation of how the area local to the site of injury was chosen. **J.** Organ-wide proliferation is decreased between resection samples from 14 to 28 dpi (p=0.0009). **K.** Local proliferation is decreased between resection samples from 14 to 28 dpi (p=0.0005). Scale bars are 100 μm.

At 28 dpi, we found that the organ-wide proliferative response was comparable to the sham-operated controls (Figure 2E-F). However, the proliferative response local to the site of injury remained significantly increased in comparison to the organ-wide response (Figure 2G). Likewise, the proliferative response local to the site of injury was still significantly increased in comparison to an analogous area in sham samples (Figure 2H-I). These results suggest that the whole-organ proliferative response had subsided at 28 dpi while the local proliferative response persisted.

We also compared the injury response between 14 and 28 dpi pancreas samples. Compared to 14 dpi, there was a significant decrease in proliferation both throughout the organ and local to the site of injury at 28 dpi (Figure 2J-K).

### Pancreatic resection induces β−cell regeneration

The identity of proliferating cells is important for understanding the regenerative mechanisms at play, which will ultimately determine whether an organ will be capable of proper function after regeneration occurs. We identified and quantified EdU positive cells expressing insulin/ proinsulin protein (from here referred to as insulin positive) in order to understand the injury response of β−cells (Figure 3A-D). We found that β−cells proliferated at a significantly higher rate in resected samples when compared to the sham, indicative of β−cell regeneration in response to injury (Figure 3E).^10^ Additionally, at 14 dpi, there was a significant increase in the number of insulin positive β−cells in the resection condition; we refer to this as increased β−cell mass (Figure 3F). By 28 dpi, there was a non-significant difference in the amount of insulin positive β−cells between the sham and resected samples (Figure 3G-H).

**Figure 3.**
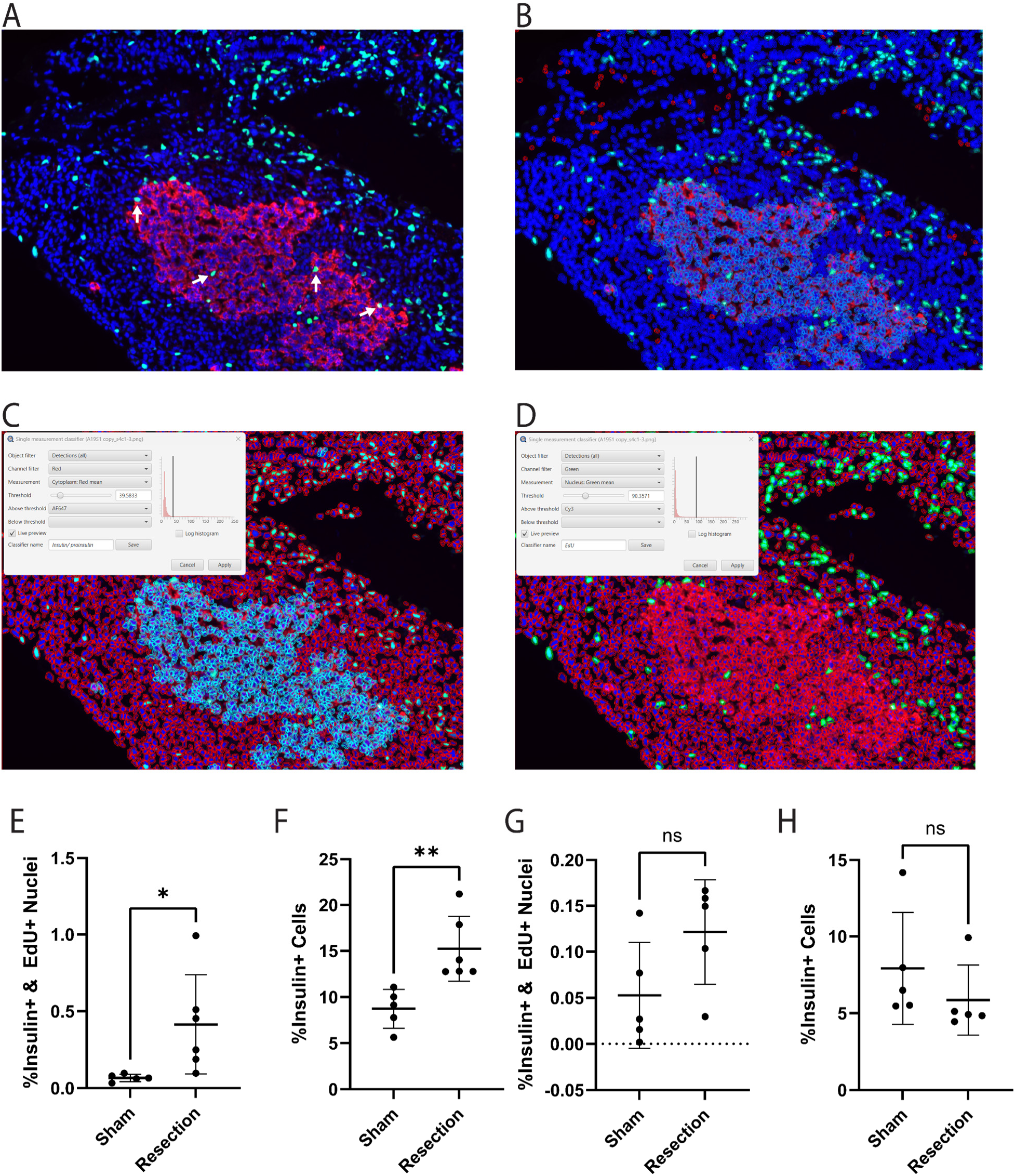
β−cell proliferation is observed at 14 days post injury but subsides by 28 days. **A.** Representative image of a coronal section of regenerating pancreatic tissue at 14 dpi. White arrows point to proliferating β−cells. **B.** Visualization of image quantification with QuPath Software. Cells were categorized by expression of insulin/ proinsulin antibody and EdU stain, then counted for downstream statistical analysis. **C.** Settings used to identify insulin/ proinsulin positive cells. Light blue outlines represent insulin/ proinsulin positive cells. Red outlines represent cells negative for insulin/ proinsulin. **D.** Settings used to identify EdU positive cells. Green outlines represent EdU positive cells. Red outlines represent EdU negative cells. **E.** A significant increase in cells positive for both EdU and insulin antibody was observed at 14 dpi (p = 0.0463) (sham n=5, resection n=6). **F.** A significant increase in insulin positive cells was observed at 14 dpi (p = 0.005) (Sham n=5, Resection n=6). **G.** No significant difference in cells positive for both EdU and insulin antibody was observed at 28 dpi (p = 0.0936) (Sham n=5, Resection n=5). **H.** No significant difference in insulin positive cells was observed at 28 dpi (p = 0.3181) (Sham n=5, Resection n=5).

### Axolotl pancreatic resection induces transcriptional changes in well-studied injury response pathways

To identify changes in gene expression associated with pancreatic resection, we generated cDNA libraries (n=4 sham, n=4 resection) with polyadenylated RNA extracted from whole-organ pancreas tissue at 14 dpi. cDNA was sequenced using the Oxford Nanopore Technologies (ONT) PromethION sequencing platform. Sequencing returned 160.34M reads, of which 138.6M reads received a qscore greater than 8. After filtering by quality, 97.79% of reads were successfully demultiplexed by their respective barcode (Figure 4A). Classified reads were aligned to the axolotl transcriptome with a median read alignment identity of 98.41% and alignment N50 of 526.48 bp (Figure 4B).^16^ Subsequent analyses of the sequencing data showed 1618 differentially expressed genes between the sham and resection groups. A heatmap and volcano plot were created in order to show several of the most significant transcriptional changes between conditions (Figure 4C-D). KEGG analysis revealed several categories to which these genes can be attributed, such as cell cycle markers, regeneration markers, antitumor markers, transcription factors, metabolic pathways, and protein transporters.

**Figure 4.**
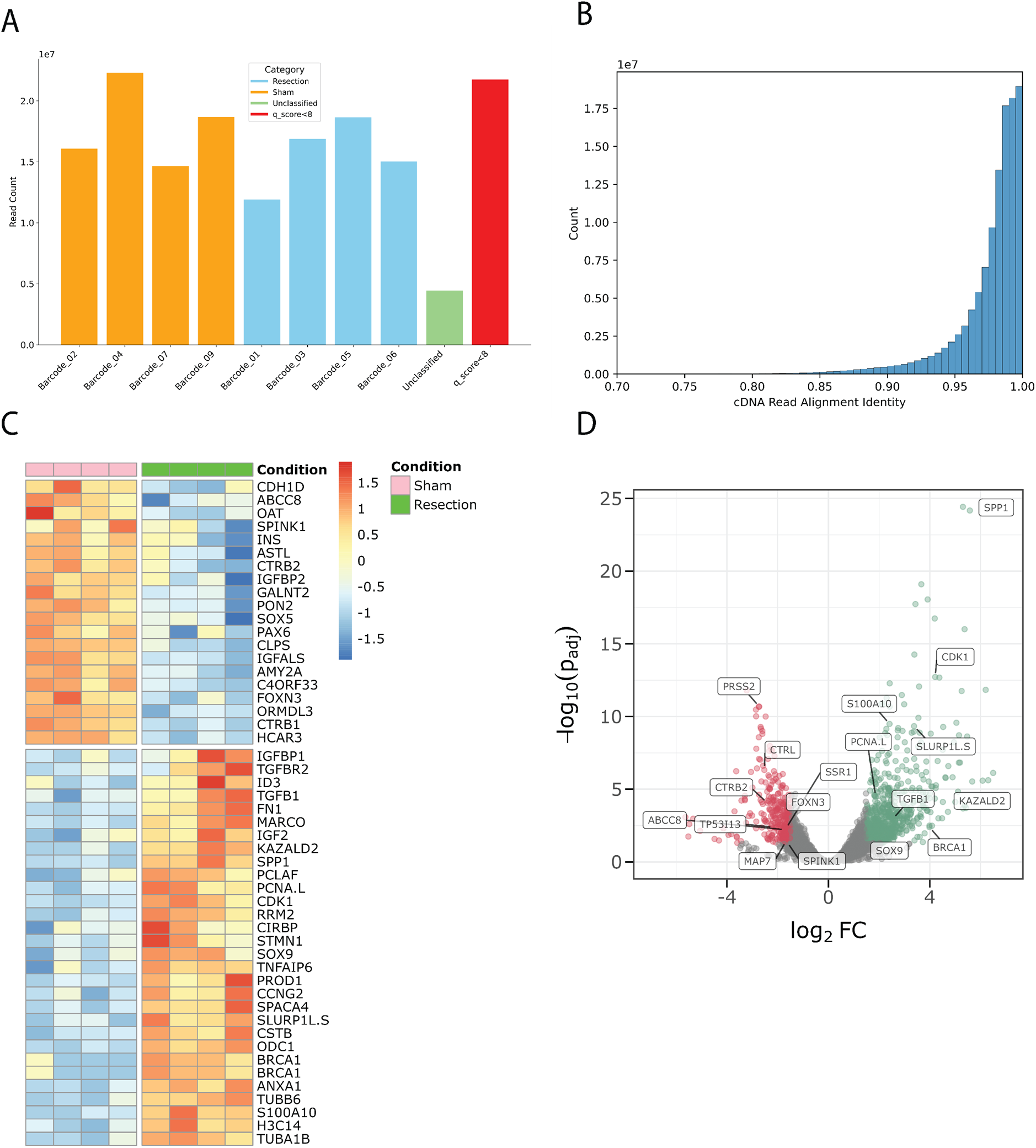
Nanopore sequencing data reveals transcriptomic changes in response to injury. **A.** Bar plot detailing reads per sample after demultiplexing. Unclassified reads and reads with a qscore less than 8 were filtered out. **B.** Histogram detailing aligned read identity vs total read counts. **C.** Heatmap of differentially expressed genes at 14 dpi **D.** Volcano plot of differentially expressed genes at 14 dpi.

Many of these genes have been implicated in axolotl limb regeneration such as *Kazald2*, which was found to be differentially upregulated.^17^ We also found that genes *Cirbp* and *Marco* were differentially upregulated, which we validated via HCR-FISH (Figure 5A-B). Interestingly, *Kazald2* and *Cirbp* are regeneration markers in other contexts such as axolotl limb regeneration.^17^ *Marco* is a macrophage marker, suggesting an immune response is provoked by pancreatic resection, which parallels the macrophage-mediated immune response necessary in axolotl limb regeneration and in a variety of other organ/appendage regeneration contexts in other species.^18,19,20^ These findings suggest that pancreatic response to injury shares characteristics with other regenerative tissues and organs in the axolotl.^17,19^ Additionally, we found *Tgf-β1* to be differentially upregulated. *Tgf-β1* is a key regulator of wound epithelium formation, blood clot formation, and inflammation in mammals, all of which are essential for regeneration to take place in axolotls.^21^ Further, Levesque *et al.* functionally demonstrated *Tgf-β1* to be essential in the initiation and control of the axolotl limb regeneration process.^21^ Parallels in axolotl limb and pancreas regeneration may reveal conserved regeneration pathways that are present in all axolotl injury responses.

**Figure 5:**
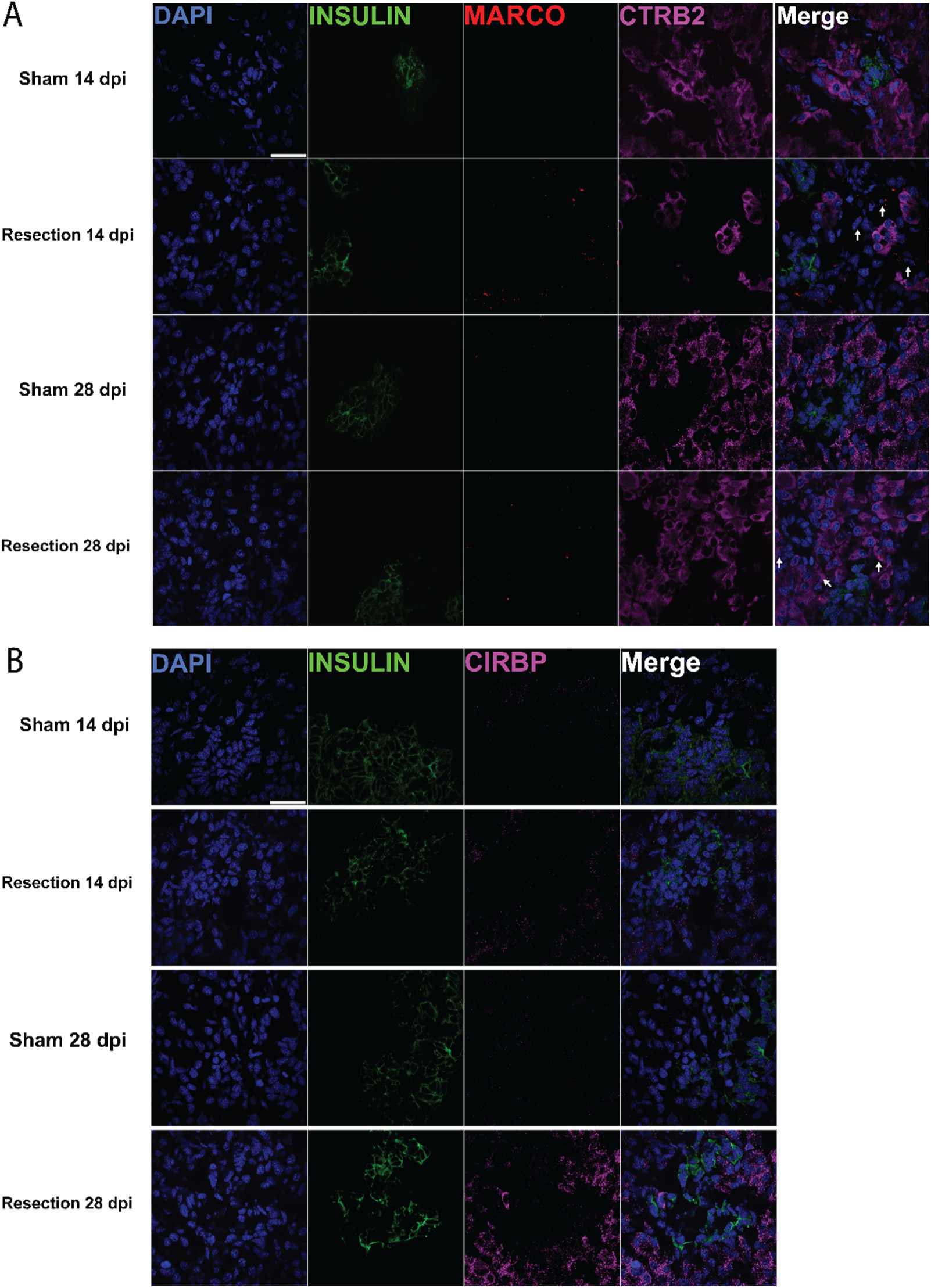
Genes previously implicated in regeneration exhibit upregulation in axolotl pancreatic injury model. **A.** HCR-FISH showing differential expression of insulin and *Marco* transcripts. *Ctrb2* was not found to be differentially expressed; however, it provides structural context for exocrine tissue. *Marco*+ cells shown with arrows. **B.** HCR-FISH showing differential expression of insulin and *Cirbp* transcripts. Scale bars are 50 μm.

We found several pancreatic development and maintenance genes to be differentially expressed such as *Ctrb2, Ins,* and *Pdx1*. *Ctrb2* is a gene encoding chymotrypsin, an essential molecule produced by the exocrine pancreas to aid in digestion. We use *Ctrb2* to visualize the distinction between the exocrine and endocrine pancreas (Figure 5A). Interestingly, both *Pdx1* and *Ins* were found to be downregulated at 14 dpi. *Ins* is the transcript responsible for insulin production by β−cells. *Pdx1* is a master regulator of pancreatic development and function. *Pdx1* is expressed throughout the gastrointestinal tract and central nervous system during development; however, it is primarily expressed in β−cells in early development and persists into maturity.^22^ Downregulation of these transcripts supports our hypothesis that β−cell regeneration has exceeded the β−cell mass needed for homeostatic function, leading to extinguished expression of transcripts responsible for β−cell development and insulin production.

### Axolotl pancreatic tissue morphology

The axolotl pancreas is a thin, triangular tissue connected directly to the base of the duodenum, with connections to the liver (Figure 1B). It has been reported that the axolotl pancreas exhibits protein level insulin expression similar to the mammalian pancreas.^23^ However, the expression of other common endocrine and exocrine pancreatic markers in axolotl is not well documented. We utilized hybridized chain reaction-fluorescent in situ hybridization (HCR-FISH) to show spatial transcript level expression of several pancreatic markers. This staining showed the presence of *Ins*-(insulin) and *Nkx6-1*-expressing β−cells, *Gcg-* (glucagon) expressing α-cells, *Sst-* (somatostatin) expressing δ-cells, (Figure 6A-B). *Pdx1*, a critical gene in pancreatic development and function, was found to be expressed in both endocrine and exocrine cells (Figure 6B).^22^ These results demonstrate that axolotl pancreatic tissue morphology shares similarities with both mice and humans, suggesting that pancreas studies in axolotl could have functional therapeutic relevance to mammalian research (Figure 6C).^8,24^

**Figure 6.**
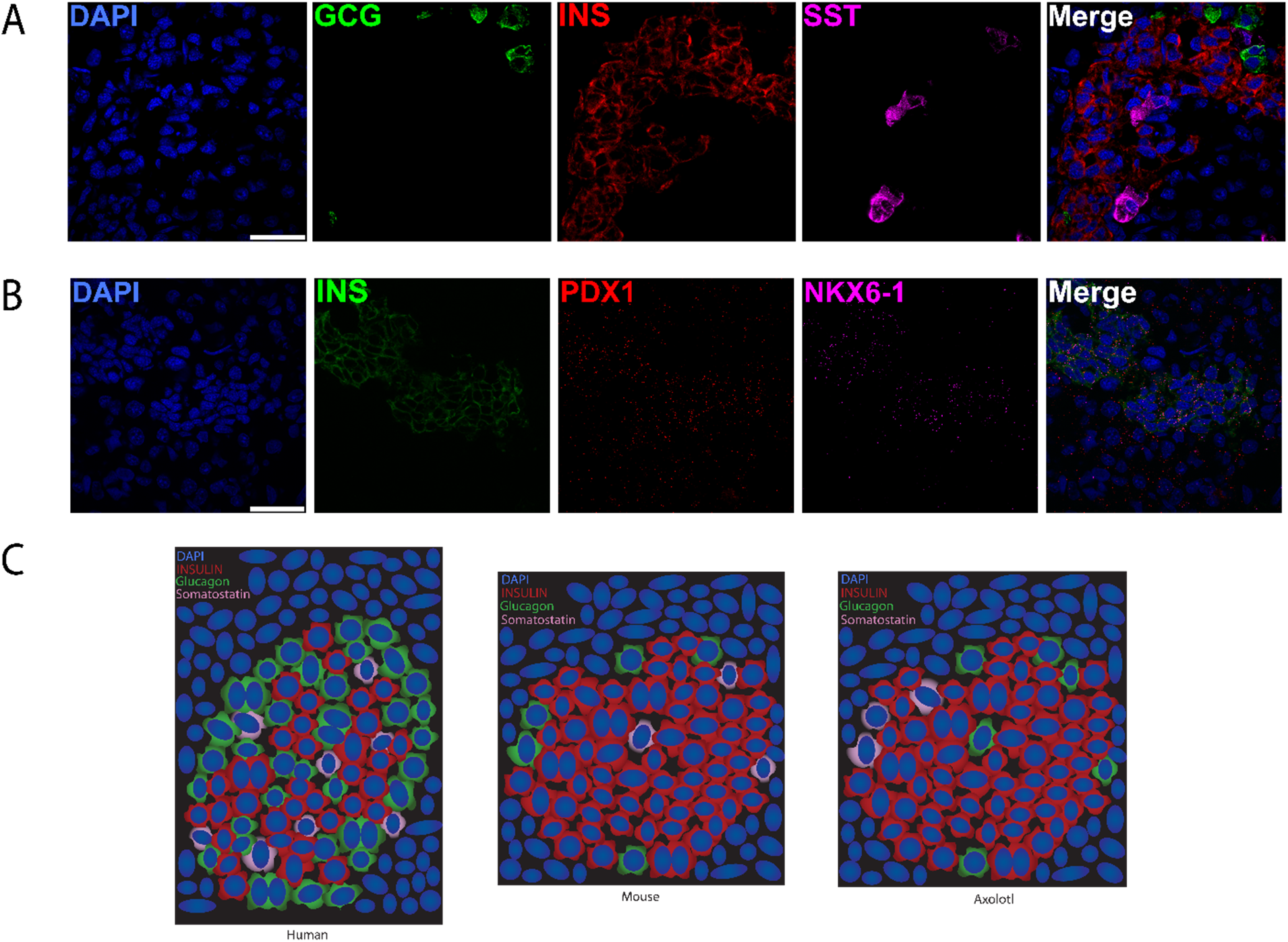
Transcript level pancreatic islet morphology in axolotl. **A.** HCR-FISH of glucagon (*Gcg*), insulin (*Ins*), and somatostatin (*Sst*) transcripts. **B.** HCR-FISH of insulin (*Ins*), *Pdx1*, and *Nkx6-1* transcripts. **C.** Cartoon representation comparing axolotl islets to mice and humans. Mouse and human islet cartoons based on data from Abdulreda *et al*^25^. Scale bars are 50 μm.

## Discussion

### Pancreatic regenerative responses are observed after injury

Axolotl limb regeneration is well documented as a blastema-mediated regeneration process. Our observations from the pancreatic resection surgery do not show any blastema-like structure—the plane of resection remains visible, and no tissue growth was visually observed at the site of injury (Figure 1B). However, non-blastema mediated regeneration has been reported in the axolotl from studies on liver regeneration via compensatory growth.^13^ Liver regeneration in the axolotl materializes in whole-organ cell proliferation, which replenishes the mass of liver removed by partial hepatectomy.^13^ At all times throughout the liver regeneration process, there is no blastemal structure, and the plane of resection remains visible.^13^

In the context of axolotl pancreas regeneration, our data suggests a similar, non-blastema mediated regeneration process. We observed a whole-organ proliferative response over the course of 14 days which showed an overall reduction by 28 days (Figure 2J-K). This is a process observed in both axolotl limb and liver regeneration; proliferation is increased until functional tissue restoration takes place, after which proliferation subsides.^13,24^ A portion of these proliferating cells were identified as β−cells and a significant increase in β−cell mass was observed at 14 dpi (Figure 2E-H). Significant increases in β−cell proliferation is indicative of β−cell regeneration in mice.^10^ Interestingly, our sequencing results indicated that transcription of insulin was downregulated at the 14 dpi timepoint. We can evoke several different possible explanations for pancreatic cells downregulating insulin transcription post-injury. For example, the regenerative response to injury may have overcompensated with more β−cell proliferation than needed, which in turn led to a downregulation in insulin production. Increased β−cell mass could mean each individual β−cell would need to produce less insulin than normal in order to maintain normal insulin levels. However, developing these hypotheses further and testing them will require additional experimentation outside the scope of this work.

### Axolotl pancreas morphology

Here, we show transcript-level expression of evolutionarily-conserved pancreatic hormones such as insulin, glucagon and somatostatin in the axolotl via HCR-FISH. Additionally, key developmental markers such as *Nkx6-1* and *Pdx1* are shown to be conserved in this context. These transcriptional characteristics establish the axolotl as a comparable model of pancreatic development and function to mice and humans.^8^

Future studies should aim to investigate the functional role of notable genes in promoting or preventing pancreatic cell function and proliferation in the context of injury. The animals used for this study were too small to tolerate repeated blood draws of the quantity needed for ELISA assays. As such, our findings would benefit from future studies that identify whether the injury-induced increase in β−cell mass in axolotl allows for the maintenance of functional insulin homeostasis. Future work should also characterize the origin of new β−cells. There is significant debate in the field as to whether pancreatic stem cells exist in mammals.^10^ Axolotl salamanders have been demonstrated to use several stem-cell-mediated mechanisms for regeneration.^20^ Lineage tracing should be performed in axolotl pancreatic injury models to ascertain whether new β−cells originate from existing β−cells or from pancreatic stem cells.

We conclude that axolotl pancreatic islets closely resemble those in mice and humans, demonstrated by the conservation of several key pancreatic gene markers (Figure 6A-B). Further, a significant increase in β−cell mass is a direct parallel between mouse and axolotl pancreatic regeneration (Figure 3F).^10^ Although mouse models are very well characterized in the context of pancreatic regeneration, translating these insights into natural β−cell regeneration has not yet manifested in humans. We assert that the axolotl is an important animal model to study because its highly regenerative nature could reveal additional factors required to improve regenerative outcomes in humans.

### Experimental Procedures

#### Pancreas resection surgery

All animal experimentation was approved by and conducted in accordance with Harvard University’s Institutional Animal Care and Use Committee (Protocol # 19-02-346). Adult wild type axolotl salamanders, 16 cm in length, were anesthetized in 0.1% tricaine solution and subjected to either a pancreatic resection surgery or a sham control surgery. The sham surgery consisted of a 2-inch lateral incision of the abdomen followed by gentle perturbation of the pancreas. The abdominal incision was closed with two horizontal mattress stitches using 6-0 nylon sutures. The resection surgery consisted of a similar 2-inch lateral incision of the abdomen followed by an approximate 20% resection of the pancreas starting from the base of the duodenum. Substantial bleeding was observed. Sterile wooden cotton tipped applicators were used to successfully clot the bleeding in all cases. The abdominal incision was closed with two horizontal mattress stitches using 6-0 nylon sutures. The animals were allowed to respond to this injury for 14 days, or 28 days post-incision, after which the whole pancreas was collected and used for histology/ sequencing.

#### Tissue collection and cryosectioning

Upon collection of the whole pancreas at 14 days, or 28 days post incision, animals were perfused with diethyl pyrocarbonate phosphate buffered saline (DEPC PBS) via aortic injection in order to clear tissue of blood and digestive enzymes while preserving RNA integrity. Tissue samples were subsequently preserved using various methods based on the intended analysis of that tissue:

Pancreas tissue samples intended for use in histology were transferred to a 4% paraformaldehyde in DEPC PBS solution upon collection. Samples were rocked overnight at 4°C. The following morning, samples were washed with DEPC PBS and transferred to a 30% sucrose in DEPC PBS solution. Samples were rocked overnight at 4°C. The following morning, samples were transferred to an OCT −30% sucrose mixture for 1 hour at room temperature. Subsequently, samples were embedded in 100% OCT on dry ice. Samples were stored at −80°C until further use. Tissue samples embedded in OCT were cryosectioned on the Leica cm 1950 cryostat at −22°C into 10 μm sections and mounted on Fisher Brand Superfrost Plus microscope slides. Samples were stored at −80°C.

Pancreas tissue samples intended for bulk RNA-seq were subject to a modified protocol based on Jun et al. which required samples to be injected with RNAlater upon collection and minced into small pieces to maximize tissue surface area contact with RNAlater^27^. Samples were subsequently transferred into 500μL of RNAlater and snap frozen on dry ice. Samples were stored at −80°C until further use. RNA extraction was completed using a phenol/chloroform phase separation with subsequent clean up using the Zymo Clean & Concentrator 5 kit.

#### EdU incorporation

A stock 5-ethynyl-2’-deoxyuridine (EdU) solution was made by dissolving EdU in dimethylsulfoxide (DMSO) at a concentration of 5μg/μL. This stock solution was diluted to 0.1μg/μL in 0.7× PBS for injection. Animals were anesthetized in 0.1% tricaine solution for 30 minutes prior to injection. EdU solution was injected intraperitoneally with care not to puncture internal organs. Each animal received 20μL per gram body weight of the injection solution. Animals were returned to normal housing conditions to incorporate with EdU for 18 hours post injection. Tissue collection immediately followed.

#### EdU and immunohistochemistry

Slides were rehydrated in 1× PBS and permeabilized with 0.5% Triton-X PBS (TX-PBS) for 30 minutes. EdU reaction was completely using copper sulfate (CuSO_4_), ascorbic acid (C_6_H_8_O_6_) and sulfo-Cy3 azide. Slides were blocked with 2% BSA in 0.1% TX-PBS for 30 minutes followed by addition of primary antibody (1:200 in blocking solution) against insulin/ proinsulin overnight (D3E7 (5B6/6), ThermoFisher Scientific, cat no. MA1-83256). Secondary antibody was added for 2 hours (AlexaFluor Donkey Anti-Mouse 647, ThermoFisher Scientific, cat no. A-31571).

#### HCR FISH

Oligonucleotides were designed using a probe generator model developed by the Monaghan Lab at Northeastern University with transcripts identified in our sequencing data through alignment to the Nowoshilow transcriptome^16^. HCR stain was completed as described by Lovely *et al.*^28^.

#### Imaging and quantification

Histological imaging was performed using the Nikon Spinning Disk Microscope, Zeiss Axioscan Microscope, and Zeiss LSM 900 Microscope. EdU & insulin/ proinsulin stains were quantified using Cell Profiler and Qupath. Technical replicates were quantified individually and then averaged into a single biological replicate. For EdU+ nuclei, all EdU+ nuclei in a section were counted and normalized to the total number of nuclei in that section. This was expressed as %EdU+ nuclei, calculated as (EdU+/total nuclei) * 100. Qupath used a spatial categorization method to count insulin/ proinsulin & EdU+ nuclei. Similarly, the proportion of insulin/proinsulin and EdU+ nuclei was determined by counting these nuclei in a section and normalizing them to the total number of nuclei in the section, expressed as %insulin & EdU+ nuclei = (insulin & EdU+/total nuclei) × 100. Generally, tissue sections were composed of 5-10 thousand total nuclei. Biological replicates at 14 dpi: Sham n=5; Resection n=6. Biological replicates at 28 dpi: Sham n=5; Resection n=5.

For selection of area local to the site of resection, a 350 x 500 μm control area was selected randomly along the plane of resection in histology images. Nuclei within the control area were analyzed as described above. A comparable area in sham samples was selected by utilizing a 350 x 500 μm control area placed along the edge of the tissue section. Nuclei in the control area were analyzed as described above. Randomized repetitions of control area placement yielded comparable results.

#### RNA extraction

Whole pancreas tissue from the sham and resection groups (n=4) was transferred to 500mL of Trizol and homogenized for 30-seconds with the Bio-Gen PRO200 Hand-Held homogenizer. A phenol chloroform phase separation was performed, and the aqueous layer was transferred to Zymo spin columns. The RNA was cleaned up following the protocol for the Zymo Clean & Concentrator 5 kit. RNA quality was determined by Nanodrop spectrophotometer, Tapestation, and Qubit.

#### Nanopore cDNA Sequencing

High quality RNA from the previous step was library prepped for sequencing according to protocol for the ONT PCR cDNA kit with barcode capability (SQK-PCB111.24). Each sample was barcoded, the libraries were combined and sequenced simultaneously on two PromethION flow cells.

Sequencing data was analyzed using a custom in-house Nextflow pipeline. This pipeline utilized Dorado to basecall, Minimap2 to align reads to the axolotl transcriptome (Nowoshilow et al., 2018), and featureCounts to quantify the data. Differential gene expression was calculated using DESeq2.

## Acknowledgements

We would like to thank The Bauer Core Facility at Harvard University and the Harvard Center for Biological Imaging (RRID:SCR_018673) for their infrastructure and support. We would like to express our gratitude to Kelly Dooling, Isaac Adatto, Kara Thornton, Damian Bernard, Brianna Blackmore, Nicholas Cardelia, Hayden Graham, Lauryn Wilson, Omenma Abengowe, Erin Anderson, Rui Qun Miao, and Vicky Yan for their assistance with animal care. We thank Doug Melton for his support of this project and his advice. We are grateful to members of the Whited Lab for their valuable advice and discussions during this study.

## Author Contributions

Conceptualization: C.J.P., A.A., J.L.W.; Methodology: C.J.P., A.A., D.D.P., A.M.S., A.R.J., N.J.L., A.D., J.L.W.; Software: H.D.S., A.A., S.J.B, C.J.P.; Validation: C.J.P., R.T.K. A.R.J.; Formal analysis: C.J.P., D.D.P., H.D.S., S.G.A., J.L.W.; Investigation: C.J.P., A.R.J.; Resources: H.D.S., M.J., J.L.W.; Data curation: C.J.P., H.D.S., M.J.; Writing – original draft: C.J.P., J.L.W.; Writing - review & editing: C.J.P., R.T.K., H.D.S., D.D.P., A.R.J., A.A., A.D., S.G.A., M.J., J.L.W.; Visualization: C.J.P.; Supervision: C.J.P., J.L.W.; Project administration: C.J.P, J.L.W.; Funding acquisition: J.L.W.

## Funding

This work was supported by the NSF-CAREER IOS-2145925 (J.L.W.) and NICHD R01HD115272 (J.L.W.), NICHD R01HD095494 (JLW), Harvard University Faculty of Arts and Sciences (JLW), the Human Frontiers Science Program Long-term Postdoctoral Fellowship #884346 (AMS), Harvard HCRP award (ARJ), Harvard Herchel Smith Undergraduate Science Research Program (RTK), the Harvard Program for Research in Science and Engineering (RTK), and ETH Zurich SEMP award (AA).

## Declaration of Interests

JLW is a co-founder of Matice Biosciences. Other authors declare no competing interests.

## References

[1] Vos, T., et al., 2020. Global burden of 369 diseases and injuries in 204 countries and territories, 1990–2019: a systematic analysis for the Global Burden of Disease Study 2019. The Lancet 396, 1204–1222.. 10.1016/s0140-6736(20)30925-9

[2] Roger, V.L., 2021. Epidemiology of Heart Failure. Circulation Research 128, 1421– 1434.. 10.1161/circresaha.121.318172

[3] Cheng, N.-C., Tai, H.-C., Chang, S.-C., Chang, C.-H., Lai, H.-S., 2015. Necrotizing fasciitis in patients with diabetes mellitus: clinical characteristics and risk factors for mortality. BMC Infectious Diseases 15.. 10.1186/s12879-015-1144-0

[4] Zhou, Q., Melton, D.A., 2018. Pancreas regeneration. Nature 557, 351–358.. 10.1038/s41586-018-0088-0

[5] Vegas, A.J., Veiseh, O., Gürtler, M., Millman, J.R., Pagliuca, F.W., Bader, A.R., Doloff, J.C., Li, J., Chen, M., Olejnik, K., Tam, H.H., Jhunjhunwala, S., Langan, E., Aresta-Dasilva, S., Gandham, S., Mcgarrigle, J.J., Bochenek, M.A., Hollister-Lock, J., Oberholzer, J., Greiner, D.L., Weir, G.C., Melton, D.A., Langer, R., Anderson, D.G., 2016. Long-term glycemic control using polymer-encapsulated human stem cell– derived beta cells in immune-competent mice. Nature Medicine 22, 306–311.. 10.1038/nm.4030

[6] Wang, S., Du, Y., Zhang, B., Meng, G., Liu, Z., Liew, S.Y., Liang, R., Zhang, Z., Cai, X., Wu, S., Gao, W., Zhuang, D., Zou, J., Huang, H., Wang, M., Wang, X., Wang, X., Liang, T., Liu, T., Gu, J., Liu, N., Wei, Y., Ding, X., Pu, Y., Zhan, Y., Luo, Y., Sun, P., Xie, S., Yang, J., Weng, Y., Zhou, C., Wang, Z., Wang, S., Deng, H., Shen, Z., 2024. Transplantation of chemically induced pluripotent stem-cell-derived islets under abdominal anterior rectus sheath in a type 1 diabetes patient. Cell 187, 6152–6164.e18.. 10.1016/j.cell.2024.09.004

[7] Wu, J., Li, T., Guo, M., Ji, J., Meng, X., Fu, T., Nie, T., Wei, T., Zhou, Y., Dong, W., Zhang, M., Shi, Y., Cheng, X., Yin, H., Mou, X., Feng, Y., Xu, X., Dong, J., He, D., Zhao, Y., Zhou, X., Wang, X., Shen, F., Wang, Y., Ding, G., Fu, Z., 2024. Treating a type 2 diabetic patient with impaired pancreatic islet function by personalized endoderm stem cell-derived islet tissue. Cell Discovery 10.. 10.1038/s41421-024-00662-3

[8] Bonner-Weir, S., Trent, D.F., Weir, G.C., 1983. Partial pancreatectomy in the rat and subsequent defect in glucose-induced insulin release.. Journal of Clinical Investigation 71, 1544–1553.. 10.1172/jci110910

[9] Moss, J.B., Koustubhan, P., Greenman, M., Parsons, M.J., Walter, I., Moss, L.G., 2009. Regeneration of the Pancreas in Adult Zebrafish. Diabetes 58, 1844–1851.. 10.2337/db08-0628

[10] Dor, Y., Brown, J., Martinez, O.I., Melton, D.A., 2004. Adult pancreatic β-cells are formed by self-duplication rather than stem-cell differentiation. Nature 429, 41–46.. 10.1038/nature02520

[11] Duméril, A. (1867). Description de diverses monstruosités observées a la ménagerie des reptiles du Muséum d’Histoire Naturelle sur les Batraciens urodèles a branchies extérieures dits Axolotls: planche V. Librairie Théodore Morgand.

[12] Jensen, T.B., Giunta, P., Schultz, N.G., Griffiths, J.M., Duerr, T.J., Kyeremateng, Y., Wong, H., Adesina, A., Monaghan, J.R., 2021. Lung injury in axolotl salamanders induces an organ-wide proliferation response. Developmental Dynamics 250, 866–879.. 10.1002/dvdy.315

[13] Ohashi, A., Saito, N., Kashimoto, R., Furukawa, S., Yamamoto, S., Satoh, A., 2021. Axolotl liver regeneration is accomplished via compensatory congestion mechanisms regulated by ERK signaling after partial hepatectomy. Developmental Dynamics 250, 838–851.. 10.1002/dvdy.262

[14] Sørensen, P., Dittrich, A., Lauridsen, H, 2023. A Novel Animal Model in Diabetes Research: Regeneration of β cells in the Axolotl Salamander? Preprint 10.20944/preprints202301.0569.v1

[15] Aguayo-Mazzucato, C., Bonner-Weir, S., 2018. Pancreatic β Cell Regeneration as a Possible Therapy for Diabetes. Cell Metabolism 27, 57–67.. 10.1016/j.cmet.2017.08.007

[16] Nowoshilow, S., Schloissnig, S., Fei, J.-F., Dahl, A., Pang, A.W.C., Pippel, M., Winkler, S., Hastie, A.R., Young, G., Roscito, J.G., Falcon, F., Knapp, D., Powell, S., Cruz, A., Cao, H., Habermann, B., Hiller, M., Tanaka, E.M., Myers, E.W., 2018. The axolotl genome and the evolution of key tissue formation regulators. Nature 554, 50–55.. 10.1038/nature25458

[17] Bryant, D. M., Johnson, K., DiTommaso, T., Tickle, T., Couger, M. B., Payzin-Dogru, D., Lee, T. J., Leigh, N. D., Kuo, T. H., Davis, F. G., Bateman, J., Bryant, S., Guzikowski, A. R., Tsai, S. L., Coyne, S., Ye, W. W., Freeman, R. M., Jr, Peshkin, L., Tabin, C. J., Regev, A., … Whited, J. L. (2017). A Tissue-Mapped Axolotl De Novo Transcriptome Enables Identification of Limb Regeneration Factors. Cell reports, 18(3), 762–776. 10.1016/j.celrep.2016.12.063

[18] Godwin, J. W., Pinto, A. R., & Rosenthal, N. A. (2013). Macrophages are required for adult salamander limb regeneration. Proceedings of the National Academy of Sciences of the United States of America, 110(23), 9415–9420. 10.1073/pnas.1300290110

[19] Godwin, J.W., Debuque, R., Salimova, E., Rosenthal, N.A., 2017. Heart regeneration in the salamander relies on macrophage-mediated control of fibroblast activation and the extracellular landscape. npj Regenerative Medicine 2.. 10.1038/s41536-017-0027-y

[20] Leigh ND, Dunlap GS, Johnson K, Mariano R, Oshiro R, Wong AY, Bryant DM, Miller BM, Ratner A, Chen A, Ye WW, Haas BJ, Whited JL. Transcriptomic landscape of the blastema niche in regenerating adult axolotl limbs at single-cell resolution. Nat Commun. 2018 Dec 4;9(1):5153. 10.1038/s41467-018-07604-0 PMID: 30514844; PMCID: PMC6279788.

[21] Lévesque, M., Gatien, S., Finnson, K., Desmeules, S., Villiard, E., Pilote, M., Philip, A., & Roy, S. (2007). Transforming growth factor: beta signaling is essential for limb regeneration in axolotls. PloS one, 2(11), e1227. 10.1371/journal.pone.0001227

[22] Ebrahim, N., Shakirova, K., Dashinimaev, E., 2022. PDX1 is the cornerstone of pancreatic β-cell functions and identity. Frontiers in Molecular Biosciences 9.. 10.3389/fmolb.2022.1091757

[23] Hansen G, Hansen B, Jorgensen P, Vogel C. Immunocytochemical localization and immunochemical characterization of an insulin-related peptide in the pancreas of the urodele amphibian, Ambystoma mexicanum. Cell and tissue research. 1989;256(3). 10.1007/BF00225598

[24] Powers AC. Architecture and Morphology of Human Pancreatic Islets. Academic Press,; 2014:257–268. 10.1016/B978-0-12-408134-5.00016-0

[25] Abdulreda, M. H., Caicedo, A., & Berggren, P. O. (2013). A NATURAL BODY WINDOW TO STUDY HUMAN PANCREATIC ISLET CELL FUNCTION AND SURVIVAL. CellR4--repair, replacement, regeneration, & reprogramming, 1(2), 111–122.

[26] Johnson K, Bateman J, DiTommaso T, Wong AY, Whited JL. Systemic cell cycle activation is induced following complex tissue injury in axolotl. Dev Biol. 2018 Jan 15;433(2):461–472. 10.1016/j.ydbio.2017.07.010 Epub 2017 Oct 31. PMID: 29111100; PMCID: PMC5750138.

[27] Jun E, Oh J, Lee S, et al. Method Optimization for Extracting High-Quality RNA From the Human Pancreas Tissue. Translational oncology. 2018;11(3):800–807. 10.1016/j.tranon.2018.04.004

[28] Lovely, A.M., Duerr, T.J., Stein, D.F., Mun, E.T., Monaghan, J.R., 2023. Hybridization Chain Reaction Fluorescence In Situ Hybridization (HCR-FISH) in Ambystoma mexicanum Tissue, in: Methods in Molecular Biology. Methods in Molecular Biology, pp. 109–122.. 10.1007/978-1-0716-2659-7_6

